# Mitigation of the effect of high light on the photosynthetic apparatus of *Rhodobacter alkalitolerans* when grown in an alkaline environment

**DOI:** 10.1101/2023.06.30.547267

**Authors:** Mohammad Yusuf Zamal, Ch Venkata Ramana, Rajagopal Subramanyam

**Affiliations:** Department of Plant Science, School of Life Sciences, University of Hyderabad, Gachibowli, Telangana- 500046 India

**Author notes:** Corresponding author Rajagopal Subramanyam Department of Plant Sciences School of Life Sciences University of Hyderabad 500 046 India.

**Keywords:** Intracytoplasmic membranes, Light-harvesting complexes, High light, Reaction center- light-harvesting complex (RC-LH1), *Rhodobacter alkalitolerans*, Supercomplexes

## Abstract

In the phototrophic alphaproteobacteria, photosynthesis is performed by pigment-protein complexes, including the light-harvesting complexes known as LH1 and LH2. The photosystem also encompasses carotenoids to assist in well-functioning of photosynthesis. Most photosynthetic bacteria are exposed to various abiotic stresses, and here, the *Rhodobacter (R.) alkalitolerans* were extracted from the alkaline pond. We report the comparative study of the photosynthetic apparatus of *R. alkalitolerans* in various light intensities in relation to this bacterium’s high pH tolerance ability. We found that as the light intensity increased, the stability of photosystem complexes decreased in normal pH (npH pH 6.8±0.05) conditions, whereas in high pH (hpH pH 8.6±0.05) acclimation was observed to high light. The content of bacteriochlorophyll *a*, absorbance spectra, and circular dichroism data shows that the integrity of photosystem complexes is less affected in hpH compared to npH conditions. LP-BN of photosystem complexes also shows that LH2 is more affected in npH than hpH, whereas RC-LH1 monomer or dimer has shown interplay between monomer and dimer in hpH although the dimer and monomer both increased in npH. The sucrose density gradient of β-DM solubilized intracytoplasmic membranes, further evidences the pattern of monomer-dimer conversion. Additionally, thin layer chromatographic separation of isolated membrane lipids shows that phosphatidylcholine (PC) levels have increased in hpH conditions which further confirms the integrity of photosystem complexes in hpH conditions. Moreover, qPCR data showed that the subunit -c of ATPase levels was overexpressed in hpH. Consequently, the P515 measurement shows that more ATP production is required in hpH, which dissipates the protons from the chromatophore lumen. This could be the reason the photosystem protein complex destabilized due to more lumen acidification. To maintain homeostasis in hpH, the antiporter NhaD expressed more than in the npH condition.

**IMPORTANCE:** *R. alkalitolerans* is an alkaline tolerant species discovered from an alkaline pond in Gujrat India. Being a photoautotrophic photosynthetic organism, it serves as a good model organism to study the photosynthetic apparatus among phototrophic alphaproteobacteria. In nature organisms not only tackle a single abiotic stress but many including temperature, light, salinity, and many other abiotic stresses. Here we investigate how two different abiotic factors light and alkaline conditions modulate the growth and photosynthetic apparatus in a phototrophic alphaproteobacterium, *R. alkalitolerans*. Our results of this study will give leads in developing alkali-tolerant algae and higher plants.

---

Photosynthesis in alphaproteobacterial is performed by a properly oriented network of densely packed pigment-protein complexes known as light-harvesting comlex2 (LH2) and light harvesting complex1 (LH1), capturing photons and transferring the energy to the reaction center (RC) (1, 2). Excited electron passes from RC to the cytochrome (Cyt) bc1 via quinone/quinol exchange at the Qb site of the RC (3,4). Light harvesting complexes comprise roughly circularly (elliptically) arranged α- helixes with bound carotenoids and bacterial chlorophyll (BChl) pigments (5). The α- helixes comprise α & β proteins forming the oligomeric light-harvesting complexes LH2 & LH1, LH1 also encompasses the reaction center (RC), making the complex RC-LH1(6). RC-LH1 are found in two forms: monomeric where the RC is surrounded by C-shaped LH1, and the dimeric form, where the S-shaped LH1 complex aggregation surrounds the two reaction centers (7,8). The LH2 and RC-LH1 complexes follow a certain pathway of electron transfer in order to generate reducing equivalents wherein the photon absorbed by the LH2 is transferred to the LH1 and from LH1 to the reaction center, where charge separation takes place between the donor and acceptor molecules. The array of these molecules includes dimer of bacterial chlorophyll (BChl), bacterial pheophytin (Bphe), and Cyt bc1 complex making a cyclic electron flow (7,9). This purple non-sulfur gram-negative bacteria house LH1-RC and LH2 by invaginating the bacterial cytoplasmic membranes leading to the formation of spherical membranous structures known as intracytoplasmic membranes (ICMs), also called chromatophores. Some of these are attached to the membrane, whereas matured IMCs are now in the cytoplasm as the development proceeds (10,11).

Members of the genus *Rhodobacter* are metabolically versatile and can grow in aerobic and anaerobic conditions. Rhodobacter has been used to study photosynthesis, hydrogen production, and poly hydroxybutyrate (PHB) production (12). *Rhodobacter (R.) alkalitolerans* grow in alkaline culture conditions, in fact, it had been discovered from an alkaline pond (13). The strategies, while grown in alkaline conditions, include (a) increased metabolic acid production through amino acid deaminases and sugar fermentation; (ii) increased ATP synthase that couples H^+^ entry to ATP generation; (iii) changes in cell surface properties, and (iv) increased expression and activity of monovalent cation/proton antiporters (14). Since *R. alkalitolerans* is a photosynthetic bacterium that utilizes photosynthesis to produce ATP, it could pave the way to understanding the relation of alkaline (high pH) tolerance and production of ATP by photosynthetic machinery, and also its own functioning in relation to alkaline conditions. Most photosynthetic bacteria grow in extreme environmental conditions. *R. alkalitolerans* is one such photosynthetic bacteria that grow in alkali conditions. Consequently, they expose to various light conditions.

Since it is a new photosynthetic bacterium, it has not been characterized by how it can cope with various stress, especially light. Thus, in the current study, we are trying to understand the effect of high light in relation to the high pH tolerance ability of *R. alkalitolerans* on the organization of photosynthetic complexes.

## RESULTS

### Growth curve and calculation of generation time

The growth curve of *R. alkalitolerans* in different light intensities in high pH and normal pH conditions was plotted (Fig. 1A). There has been an effect of high pH on the attainment of stationary phase. High pH (hpH) grown cells entered the stationary phase around the optical density of 1.6, which took 40 h in high light (250 & 500µmol photons m^-2^s^-1^) whereas at 30 µmol photons m^-2^s^-1^it took nearly 60 h to enter the stationary phase. The normal pH (npH) grown cells took 1.9 optical density, which took 40 h to enter the stationary phase. The effect of light on generation time comes down as light intensity has increased (Table .1).

**FIG 1.**
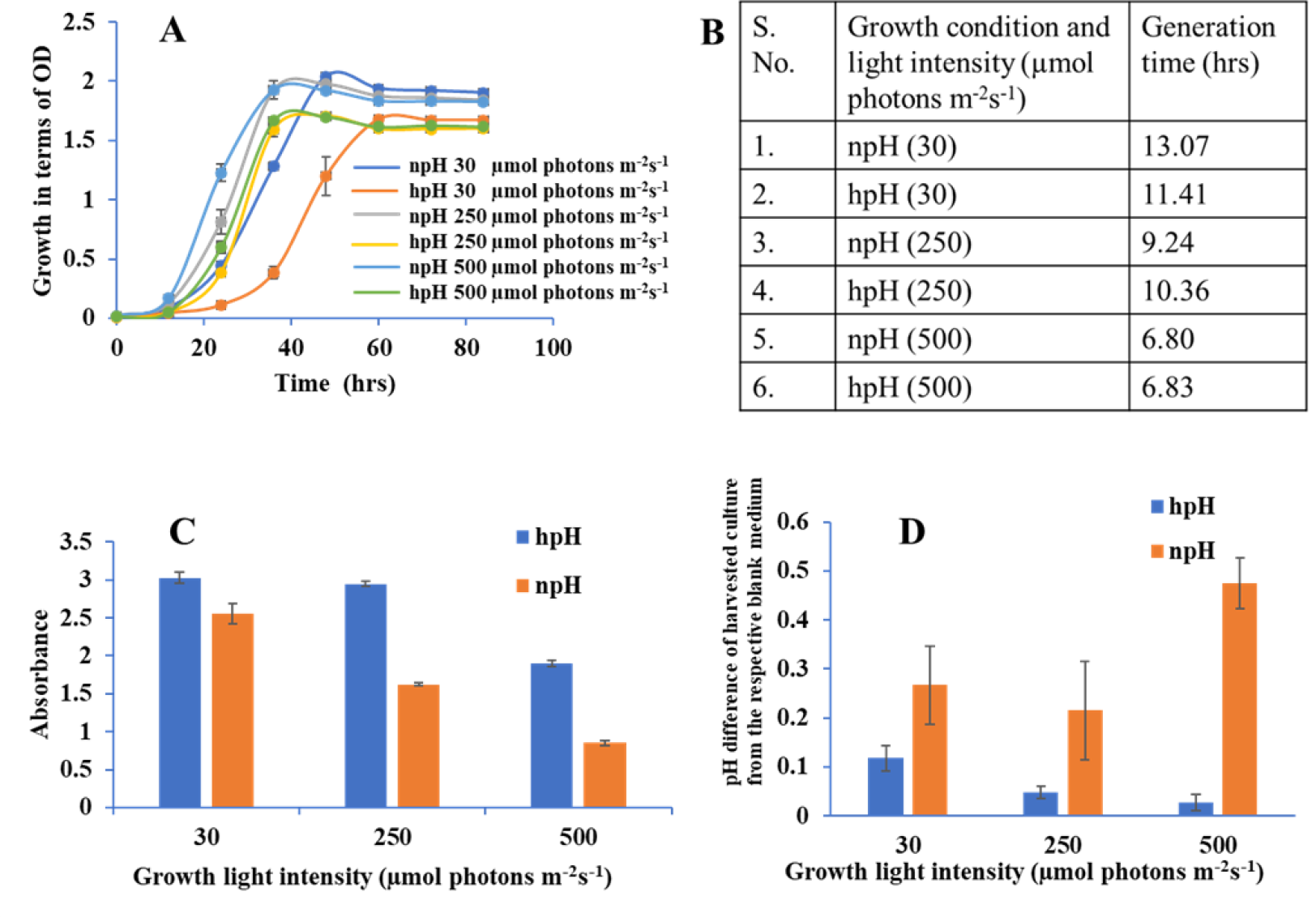
(A) Growth curve of *R. alkalitolerans* in alkaline (pH 8.6, hpH) and at normal pH (pH 6.8) in light intensities of 30 µmol photons m^-2^s^-1^, 250 µmol photons m^-2^s^-1^and 500 µmol photons m^-2^s^-1^. Cultures were grown anaerobically in 8 ml screw-capped tubes at room temperature (25℃). (B) Table .1 Generation time: Calculation of generation time/doubling time at 30 µmol photons m^-2^s^-1^, 250 µmol photons m^-2^s^-1^and 500 µmol photons m^-2^s^-1^in npH and hpH culture media. (C) Comparative representation of content of BChl *a* in hpH and npH conditions in all the three light conditions (30 µmol photons m^-2^s^-1^, 250 µmol photons m^-2^s^-1^, 500 µmol photons m^-2^s^-1^). (D) Measurement of pH difference of the culture from that of blank media in hpH and npH Culture grown at light intensity of 30, 250 and 500 µmol photons m^-2^s^-1^.

### Bacteriochlorophyll *a* estimation and carotenoid to BChl *a* ratio

Bacteriochlorophyll *a* is one of the major pigments, along with carotenoids play an essential role in bacterial photosynthesis contributing in assembly and structural stability of photosystem complexes. It is the major component of LH1-RC and LH2 and a special pair of RC. *R. alkalitolerans* photosystem also contains the pigment spheroidene which plays a very important role in photoprotection and energy transfer being the accessory light-harvesting pigment. Carotenoids have been shown to scavenge the ROS to safeguard the photosystem complexes as well during the photo-oxidative stress as a result of increased light stress (15). The content of BChl *a* has increased in a culture grown in a high pH condition compared to a normal pH condition (Fig.2), indicating that *R. alkalitolerans* accumulate more bacteriochlorophyll in hpH conditions. As carotenoids play an important role in scavenging ROS whereas BChl *a* majorly absorbs the energy, we calculated the carotenoid to BChl *a* ratio as a function of the state of the cell under photooxidative stress. Here we found that the ratio has increased with an increase in light intensity but it is more in npH than hpH (S2).

**FIG 2.**
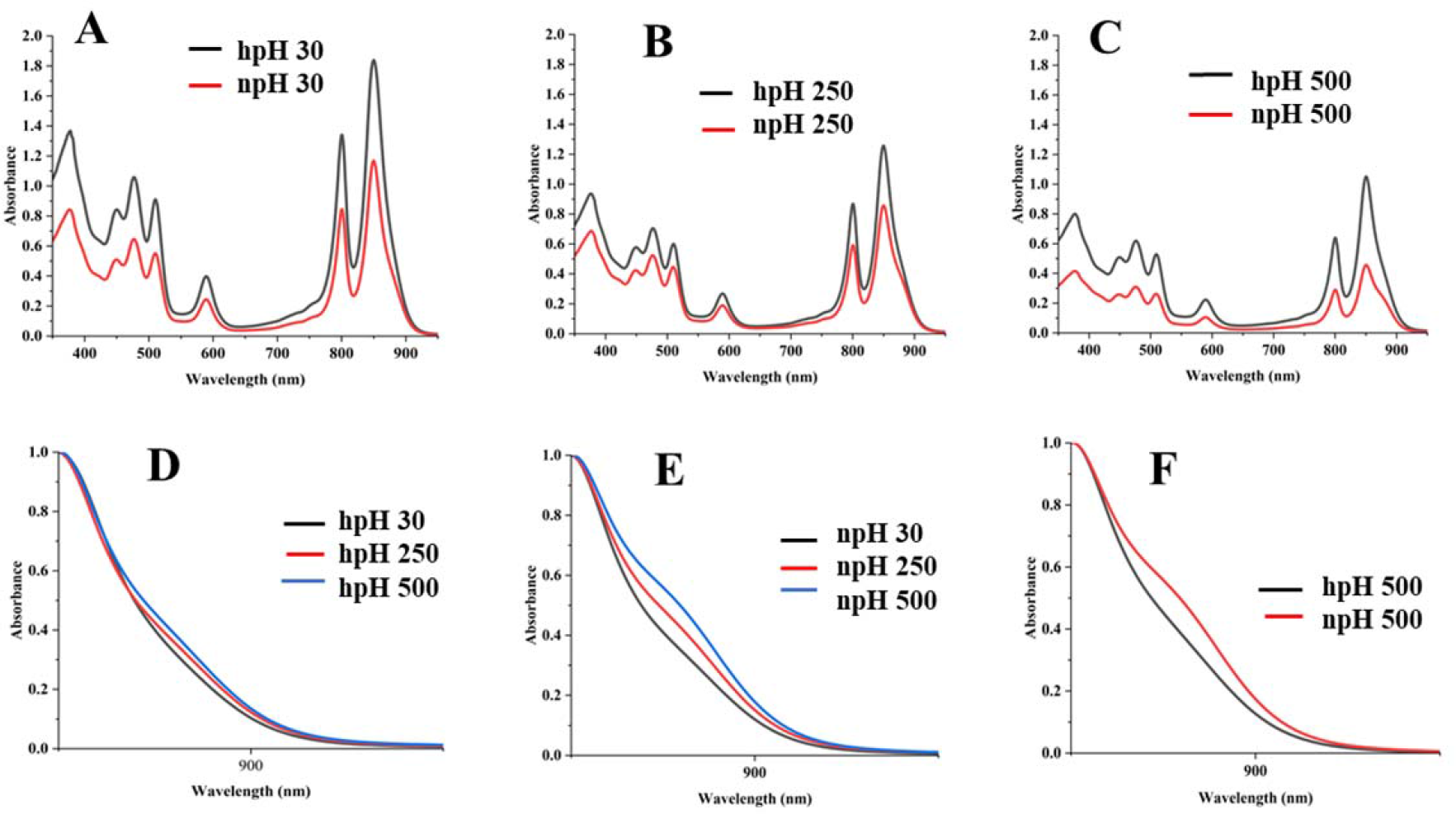
Comparative representation of absorbance spectrum of ICMs isolated from hpH and npH grown cells. (A) absorbance spectra of npH and hpH from 30 µmol photons m^-2^s^-1^. (B) absorbance spectra of npH and hpH from µmol photons m^-2^s^-1^. (C) absorbance spectra of npH and hpH from 500 µmol photons m^-2^s^-1^. (D) Enlarged view of shoulder peak of hpH ICM absorbance. (E) Enlarged view of shoulder peak of npH ICM absorbance. (F) Enlarged comparative view of absorbance the npH and hpH ICM at 500 µmol photons m^-2^s^-1^ light intensity.

### Measurement of pH of culture media while harvesting

Here we have plotted the pH difference between the normal pH harvested culture and the blank and the same with the high pH harvested culture from its blank. The pH of culture in normal pH media has increased as the light intensity increases with bit exception at 250 µmol photons m^-2^s^-1^ (Fig.1C). In contrast, in high pH grown media, the pH difference in harvested culture from that of the blank has subsequently decreased. This indicates that growing the culture at high pH leads to expression of antiporters (14). which helps in maintaining the cytoplasmic pH of cell, and photosystem integrity (Fig. 4). At the same time not let the extracellular pH increase more as compared to that of the normal pH harvested culture (grown culture). The less increase in high pH grown culture compared to blank in also might help maintain the integrity of photosystem complexes showed in BN-PAGE and circular dichroism which could be involved in maintaining the homeostasis of the cell. From this preliminary data, it has to be found out that the expression of antiporters might be helping in the dissipation of excess proton gradient built during high light in high pH culture, which leads to photoprotection in high light compared to that of the high light normal pH grown cells (14).

### Absorbance spectroscopy analysis

Comparative study of the absorbance spectrum of ICMs shows that in hpH (high pH, pH 8.6±.05), the absorbance intensity is more in all light conditions compared to npH (normal pH, pH 6.8±0.05) (Fig. 2). Absorbance spectrum is also an indication of photosystem integrity and stability of photosystem in relation to given stress condition. When increasing light intensity, the ICMs in both normal and high pH complexes are reduced, but in npH, the complexes are more sensitive than hpH. Absorbance data at equal protein concentration shows that the spectrum intensity is comparatively higher in high pH conditions compared to normal pH conditions, indicating the role of high pH tolerance ability in photoprotection. The absorbance spectra of only the RC-LH1 complex show that in hpH its peak intensity is less compared to that of the npH ICMs which signifies that in npH condition the RC-LH1 has increased with an increase in light intensity. But which form of RC-LH1 increased either dimeric or monomeric is not clear yet. This will be addressed from BN-PAGE and SDG data (Fig. 2f).

### Circular Dichroism (CD) analysis

It is important to investigate the basic arrangement of the BChls in the complex in all the respective growth conditions. CD spectrum of ICMs from all these six conditions can provide detailed information about the organization and optically active pigments. In the CD spectrum, it is obvious that organizational pattern pigment protein interaction remains the same. In contrast, the intensity of CD peaks is much stronger in hpH conditions than in npH-grown cells. The near-infrared (NIR), Q_Y_ region shows the doublet band with a strong positive band at 850 nm and a strong negative peak at 860 nm pertaining to LH2 and LH1-RC (30,16) (Fig.3). CD spectra have also shown the absorption peaks for the Q_x_ region at 590 nm and in the carotenoid region as well from 400-550 nm. Like the absorption spectra results, the CD spectra data also depicts the changes in pigment-pigment/protein complexes during an increase in light intensity. These changes are much lesser in hpH than in npH, indicating that the complexes are stabilized in high pH. Also, under high pH and high light, these complexes are stable.

**FIG 3.**
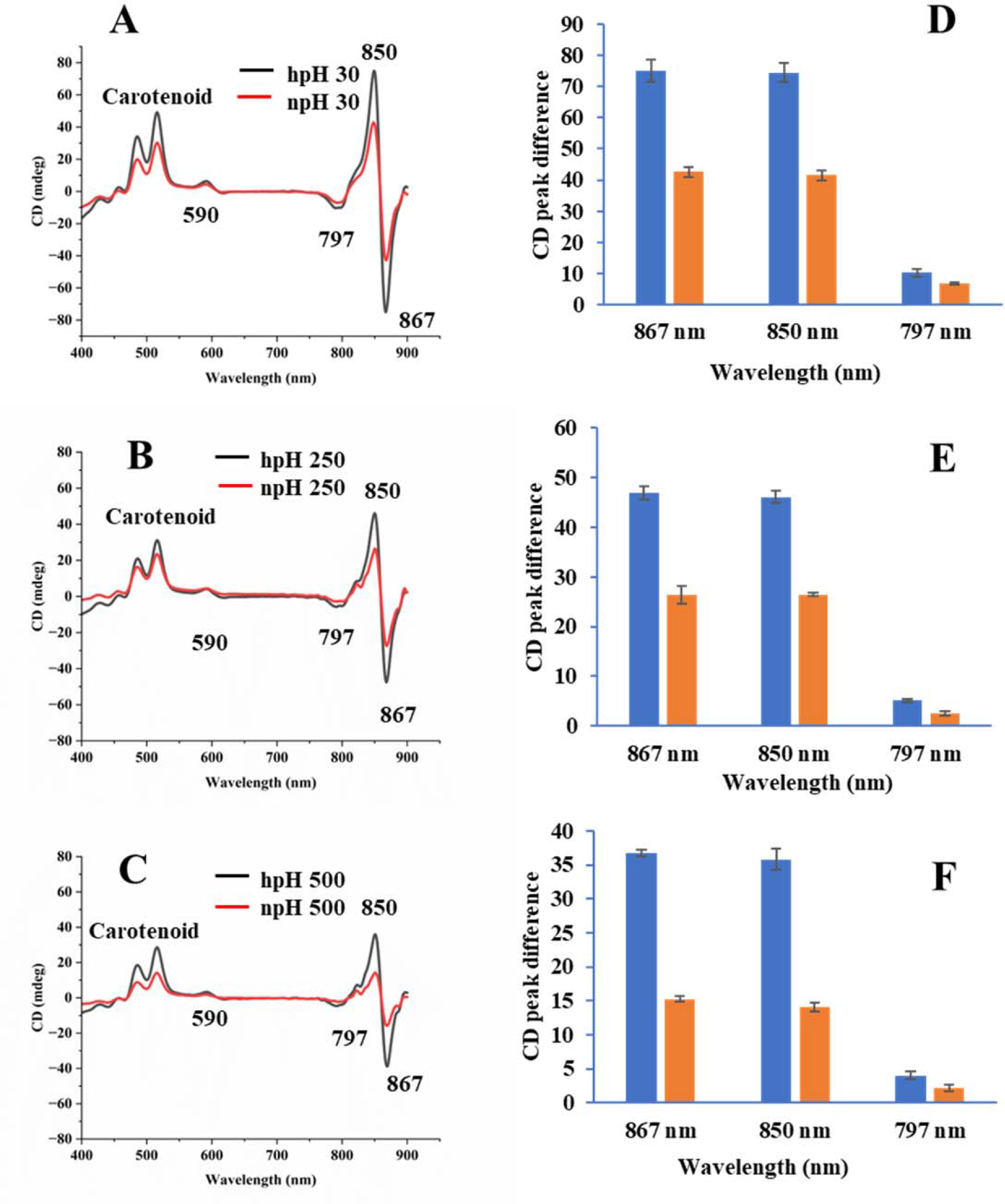
Comparative representation of circular dichroism of ICMs isolated from hpH and npH grown cells. (A) Circular dichroism spectra of npH and hpH from 30 µmol photons m^-2^s^-1^. (B) circular dichroism spectra of npH and hpH from 250 µmol photons m^-2^s^-1.^ (C) circular dichroism spectra of npH and hpH from 500 µmol photons m^-2^s^-1^. Major peak intensity differences of circular dichroism spectra of ICMs from npH and hpH in all three light conditions (D E F).

### Large pore blue native page (LP-BN) and identification of protein subunit from selected protein complex

Blue native page separates the photosystem complexes without denaturing the photosynthetic pigment-protein complexes. Here we found that as the light intensity increases, it affects the photosystem complexes. A prominent decrease is seen in the case of RC-LH1 dimer and an increase in RC-LH1 monomer in hpH condition, which could be because of the conversion of dimer to monomer. In contrast, in npH-grown cells, it is inverse of the hpH wherein both dimeric and monomeric reaction centers have increased with the increase in light intensity. This could be an interesting phenomenon that could also pave the way to understanding the real reason for dimer monomer interconversion. LH2 major and minor both have decreased with light intensity, but the extent is less in hpH conditions than in npH conditions (Fig. 4). Even in normal growth conditions, the alkaline pH shows a strong band of supercomplexes than the normal pH. The banding pattern is in accordance with the previous work (17).

**FIG 4.**
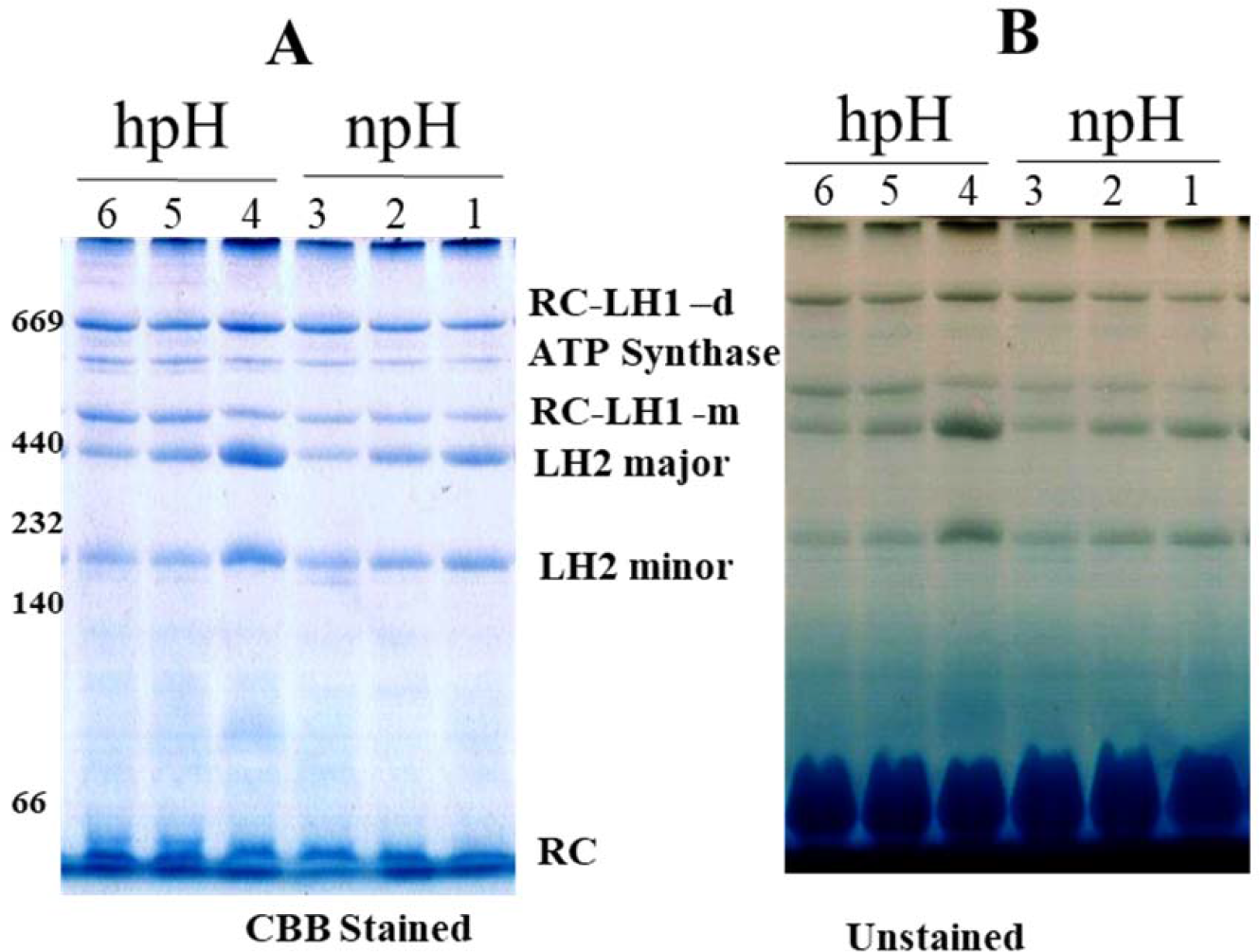
Large pore blue native page (LP-BN) of photosynthetic complexes solubilized in 1% n dodecyl-beta-d maltoside. Lane 1,2,3 represents the photosystem complexes from npH grown at light intensity of 30, 250, 500 µmol photons m^-2^s^-1^. Lane 4,5,6 represent photosystem complexes from hpH grown at a light intensity of 30,250,500 µmol photons m^-2^s^-1^. a respective growth condition CBB stained with native marker lane (A) without stain (B).

After staining the gel with colloidal Coomassie brilliant blue G -250, a non-pigmented band appeared below RC-LH1 dimeric band, which increased with an increase in light intensity in both npH and hpH conditions (Fig. S1). In order to identify this particular protein complex band second dimension of the LP-BN strip was performed. The upper subunit protein bands were identified by orbitrap high-resolution liquid chromatography-mass spectrometry (OHRLCMS). Out of these, four protein bands were identified to be ATP synthase α (55.12kDa), β(50.45 kDa), γ (31.2 kDa), and δ (19.34 kDa) (Fig. 6 B). The protein complex is found to be as ATP synthase complex increased with increase in light intensity and its expression is more in hpH condition. As light intensity increases, photons are absorbed, and the chromatophore lumen becomes more acidified because of proton accumulation. The acidic luminal condition of chromatophore leads to protein instability which could be more in npH as pumping of the proton is less in npH because the level of ATP synthase complex is less in this condition. SDS-PAGE of ICMs from each condition shows that the protein abundance of photosystem complex proteins is higher in hpH than that of npH (Fig. S1).

### Transmission Electron Microscopy

Electron micrograph of ICMs from *R. alkalitolerans* cells in all three light conditions in relation to culture condition npH shows that high light 250 µmol photons m^-2^s^-1^and 500 µmol photons m^-2^s^-1^has drastically affected the ICMs compared to that of the hpH grown cells. Looking at images from 250 and 500 µmol photons m^-2^s^-1^ of light intensity, it is quite obvious that npH- grown cells are much more affected than that of hpH grown cells, and the number of ICMs has also come down (Fig. 5). Another phenomenon is also apparent in hpH grown cells that the cell size is increased. In *E. coli* where cells grown in alkaline conditions were increased in size and length compared to cells grown at neutral pH. In our study also, the cell size has increased in high pH grown cells compared to that of the npH-grown cells. This property of elongation in cell size is more visible in normal light and whereas with an increase in light intensity, the cell size has also decreased.

**FIG 5.**
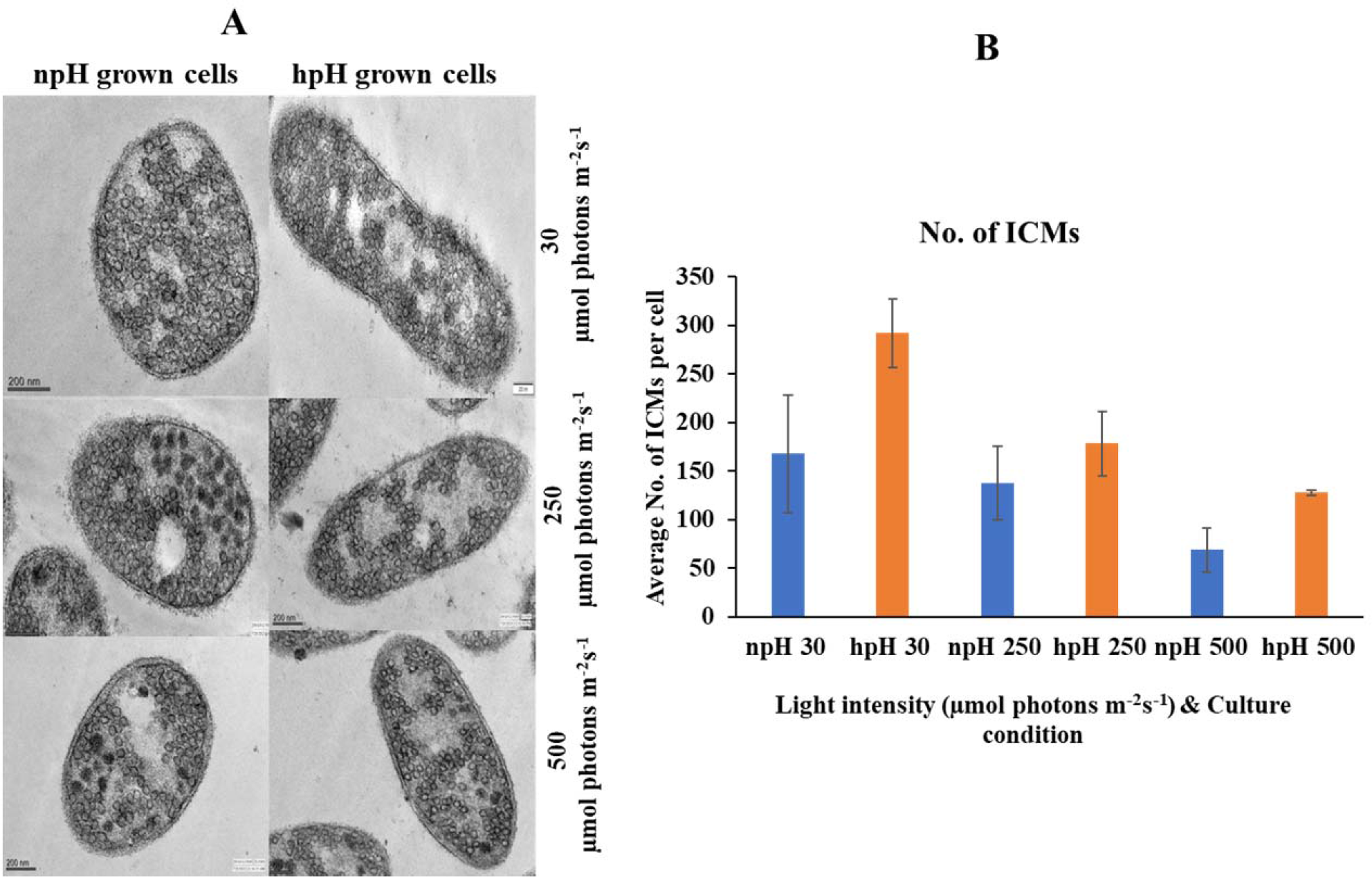
(A) Transmission electron micrograph of thin sections of cells in various light intensities in relation to pH of *R. alkalitolerans* culture condition. (B) Number of chromatophore/ ICMs from npH and hpH condition at light intensity of 30, 250, 500 µmol photons m^-2^s^-1^.

### Sucrose density gradient sedimentation and absorbance spectra of fractions from npH

Sucrose density gradient fractions were carefully collected, and their absorbance spectra were carried out. SDG shows that the F1 fraction, originating from light harvesting (LH2) has decreased with increasing light intensity. Still, the F2 fraction, which is for the monomeric reaction centre has increased with the light intensity, which may be because of a decrease in the F3 fraction, which represents the dimeric RC-LH1 complex (Fig. 6). Same is visible from the absorbance spectroscopy of each fraction of SDG. With increase in light intensity, the RC-LH1 monomer and dimer have increased as the shoulder peak at 875 nm has enhanced while increase in light intensity. It is also in accordance with the absorbance spectroscopy of the ICMs that the shoulder peak at 875 nm has increased (Fig. 2 e, f).

**FIG 6.**
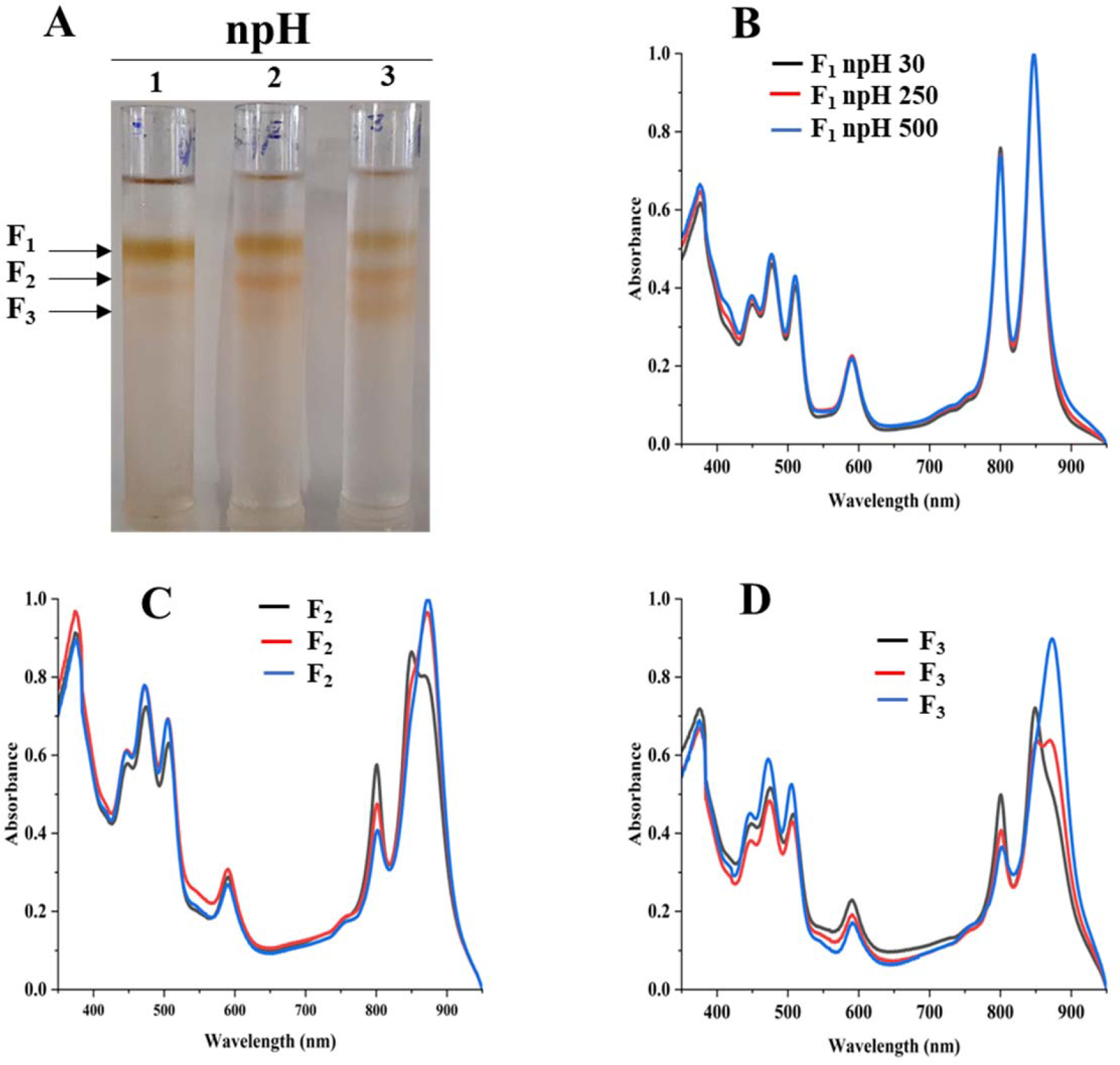
Sucrose density gradient of ICMs solubilized in 1% n-dodecyl beta-D maltoside from npH. 1,2,3 represent samples from light intensity of 30,250,500 µmol photons m^-2^s^-1^. (A). Absorbance spectra of all the three-fractions obtained F1 (B), F2 (C), F3 (D) in all three light conditions of 30,250,500 µmol photons m^-2^s^-1^.

### Sucrose density gradient sedimentation and absorbance spectra of fractions from hpH

Sucrose density gradient fractions were carefully collected, and their absorbance spectra were measured. SDG shows that the F1 fraction, which is for light harvesting (LH2) has decreased with increasing light intensity. But the F2 fraction, which is for the monomeric reaction center, has increased with light intensity. This could be because of the decrease in the F3 fraction which represents the dimeric reaction center complex, and its content was decreased (Fig. 7). It is also in agreement with Blue native PAGE (Fig. 4). The absorbance spectroscopy of each fraction further confirms that the dimeric reaction center has converted to the monomeric reaction center (Fig. 7).

**FIG 7.**
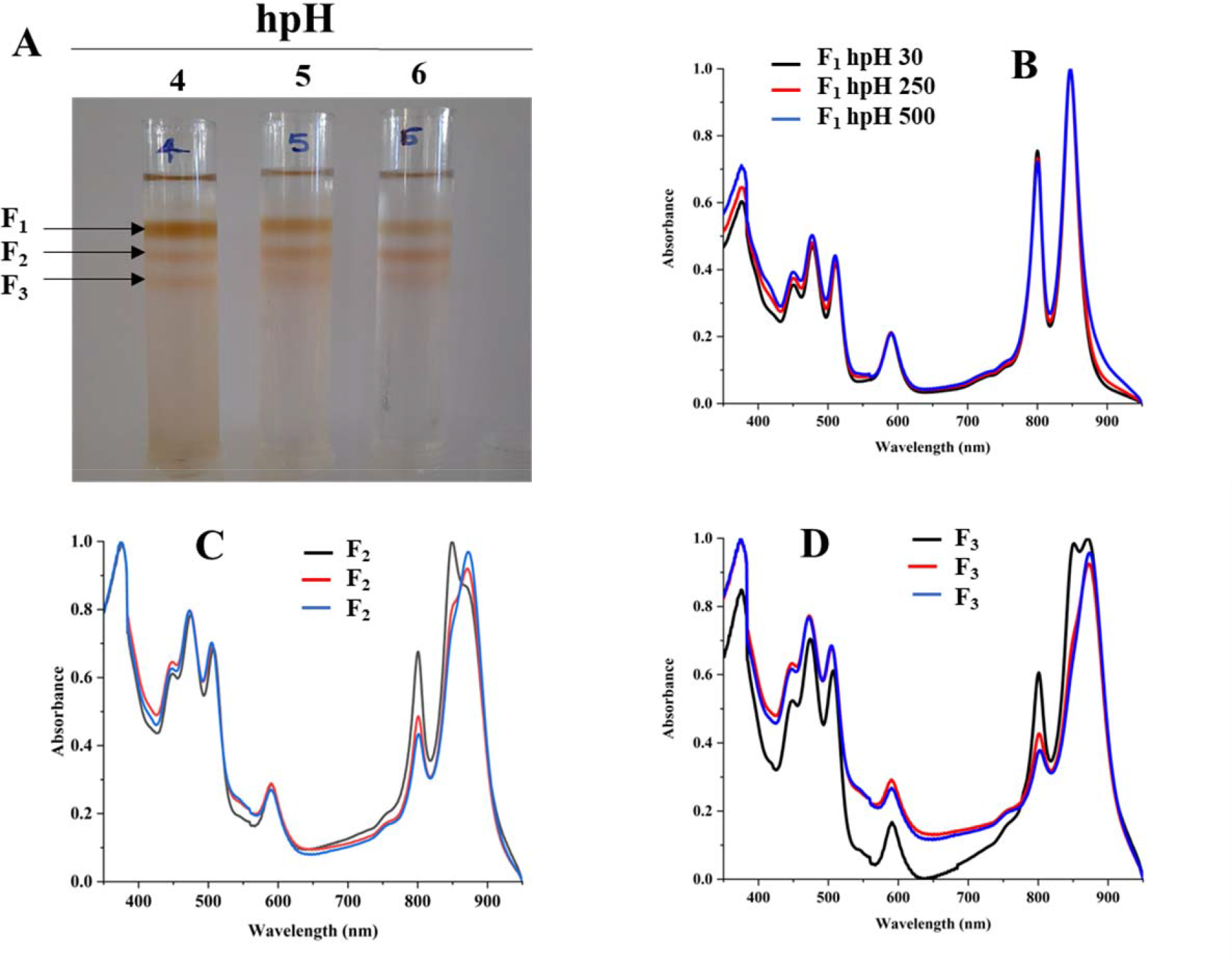
Sucrose density gradient of ICMs solubilized in 1% n-dodecyl beta-D maltoside from hpH. 4,5,6 represent samples from light intensity of 30,250,500 µmol photons m^-2^s^-1^(A). Absorbance spectra of all the three-fractions obtained F1 (B), F2 (C), F3 (D) in all three light conditions of 30,250,500 µmol photons m^-2^s^-1^.

### Quantitative real-time PCR and polar lipids

To assess the transcript level of the ATPase and the antiporter NhaD qPCR was performed for subunit ‘c’ of ATPase. Both proteins play a crucial role in the dissipation of the proton gradient. NhaD has been shown to play a significant role in the homeostasis of bacterial cells when shifted in culture media of different alkalinity (14). ATP synthase also plays a very similar role to that of the NhaD, but it utilizes ADP (adenosine diphosphate and inorganic phosphate) to make ATP which is used as an energy source in various metabolic processes in the cell. In this result, it is evident that the transcript level of both genes has increased significantly in hpH-grown cells compared to that of the npH-grown cells (Fig. 8 b & c). Moreover, both genes’ expression levels have increased in response to increase in light intensity.

**FIG 8.**
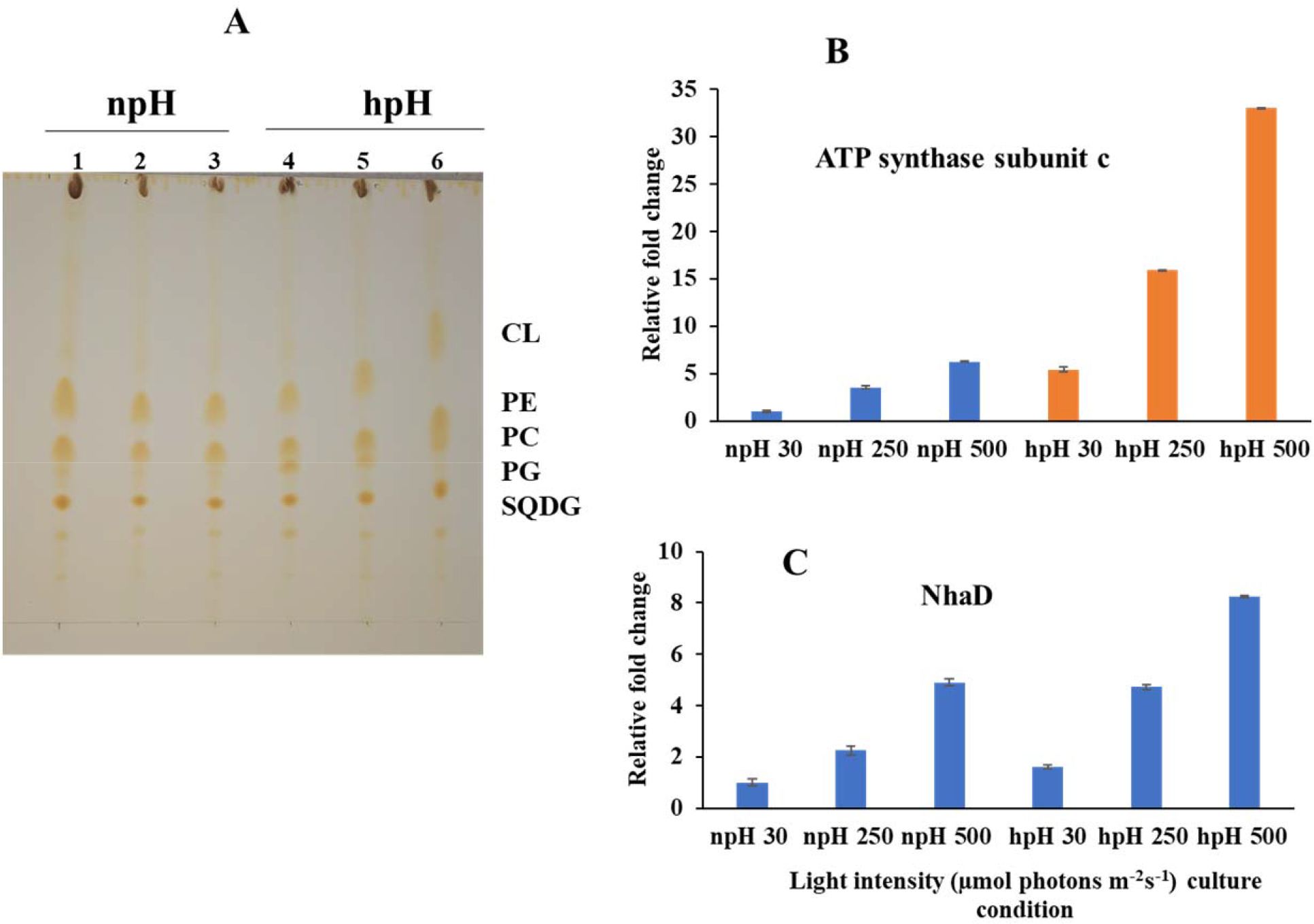
(A) Thin layer chromatographic separation of isolated polar lipids; Cardiolipin (CL) Phosphatidylethanolamine (PE) phosphatidylcholine (PC) phosphatidylglycerol (PG) and sufoquinovocyl diacylglycerol (SQDG) in npH (1,2,3 represent light intensity of 30,250,500, in all three light conditions of 30,250,500 µmol photons m^-2^s^-1^) and hpH (4,5,6 represent the light intensity of 30,250,500, in all three light conditions of 30,250,500 µmol photons m^-2^s^-1^). (B) q-PCR of subunit -c of ATPase in npH and hpH at light intensity of 30,250,500 µmol photons m^-2^s^-1^. (C) q-PCR of sodium proton antiporter NhaD in npH and hpH at light intensity of 30,250,500 µmol photons m^-2^s^-1^.

Thin layer chromatography of polar lipids isolated from complete cells shows that this bacterium has sufoquinovocyl diacylglycerol (SQDG) phosphatidylcholine (PC) phosphatidylglycerol (PG) phosphatidylethanolamine (PE) and cardiolipin (CL) as major polar lipids making lipid bilayer (Fig. 8a). After analysing the separation of lipids, it is clear that the hpH-grown cells have a higher amount of phosphatidylcholine (PC) than npH in all light intensities. Since SQDG, PC, PG, PE, and CL are proven to be involved in the stabilization of the reaction center and especially PC has been shown to fasten the electron transfer rates in charge recombination states of Q_A_ and Q_B_ (18). It supports the earlier experiment that the photosystem complexes are more stable in hpH condition than that of the npH-grown cells.

### Electrochromic shift (ECS) analysis of whole cells

P515 measurement measures chromatophore lumen acidification in relation to ATPase activity. Fig. 9 shows the electrochromic shift measurement by P515 signal measurement of intact *R. alkalitolerans* cells grown in different conditions. Sphaeroidene is the only carotenoid present in *R. alkalitolerans* cells. The electrochromic shift in carotenoid sphaeroidene and sphroidenone results from increased membrane potential due to more chromatophore lumen acidification (34,19). The relaxation of the P515 normalized signal is also an indicator of ATPase activity. In high pH conditions, the relaxation is quite faster than that of the npH cells and has increased with an increase in light intensity. Although in npH cells, the relaxation time has decreased, it is comparatively lesser than that of the hpH cells. The qPCR of the ‘c’ subunit of ATPase expression level has also increased with an increase in light intensity which is more in hpH-grown cells (Fig. 8c). It further confirms from LP-BN-2D and protein identification level of ATP synthase is more in hpH-grown cells (Fig. S2). This helps in less acidification in chromatophore lumen and less detrimental effect to photosystem complexes.

**FIG 9.**
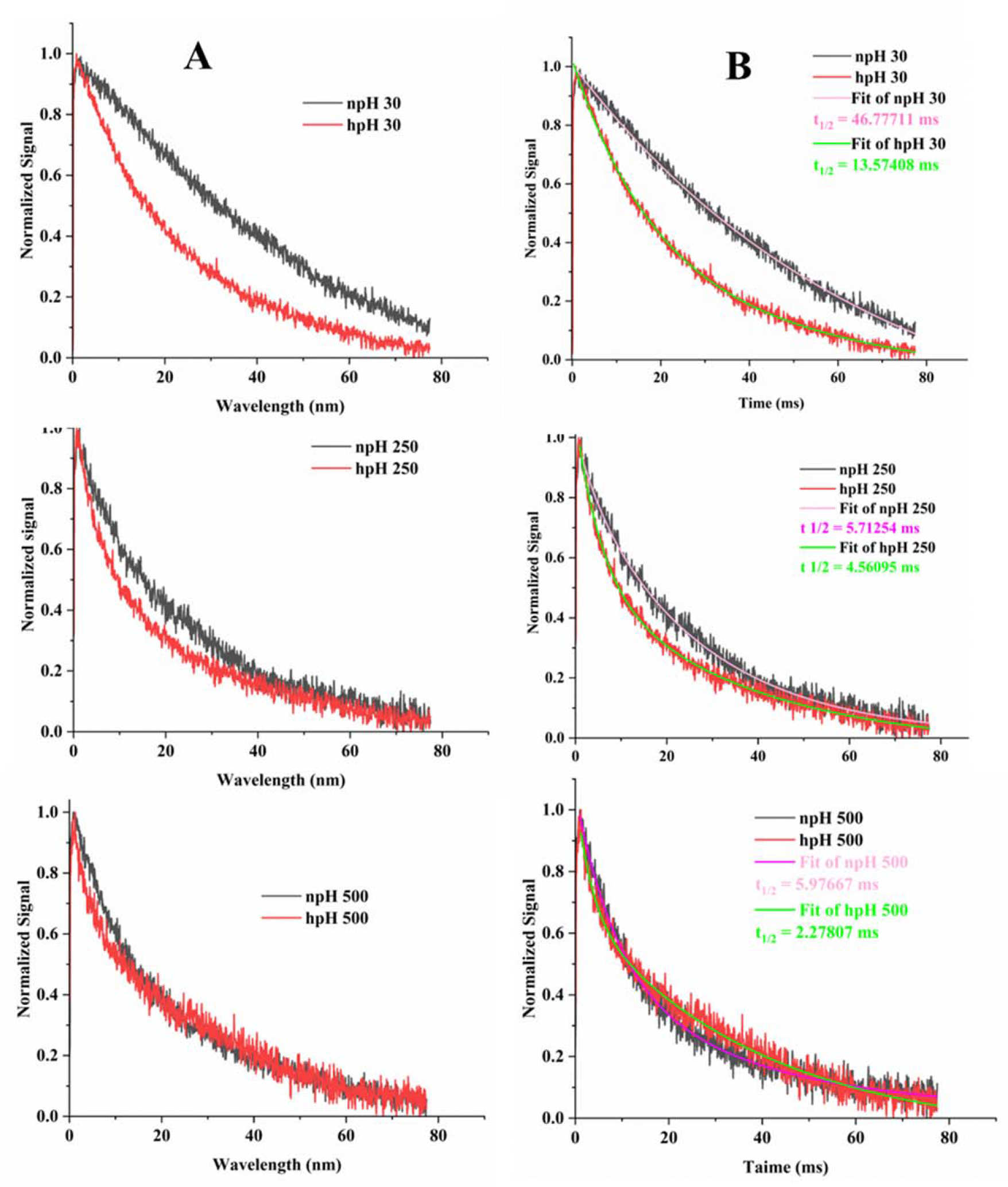
(Panel A) Flash induced absorption changes in cells of *Rba. alakalitolerans* measured at 515 nm walength. Cells were adopted for 1 hr dark incubation before the measuremnt. (Panel b) Calculation of t _1/2_ for each light intensity. Half life of each decay kinetics was calculated by double fitting in origin software. Half life (t_1/2_) for npH and hpH at 30 µmol photons m^-2^s^-1^.t_1/2=_ 46.771 ms, t_1/2=_13.57 ms. For 250 µmol photons m^-2^s^-1^. npH and hpH t_1/2=_ 5.712ms, t_1/2=_ 4.560 ms and for 500 µmol photons m^-2^s^-1^.in npH and hpH half-life were t_1/2=_ 5.976 ms, t_1/2_ = 2.27807 ms respectively.

### Estimation of total ROS and quantification of SOD

To measure the stress level in the cell in all conditions total ROS was measured by H_2_DCFDA fluorescent dye. ROS level has increased with increase in light intensity in both npH and hpH conditions. It was found that level of ROS is relatively high in npH grown cell in all three light conditions compared to in hpH condition. To further look into level of superoxide dismutase expressed in the cell, the after-sonication cell supernatant was used to quantify the level of SOD. A 20 µg of protein was loaded in non-denaturing polyacrylamide gel. Only one band of SOD appeared after staining the gel with staining solution containing Nitroblutetrazolium chloride (NBT). According to (20) it a CuZnSOD that is expressed and its expression level has increased with an increase in light intensity in both npH and hpH conditions but it is relatively less in hpH conditions.

## DISCUSSION

This study attempts to understand the photosynthetic complexes of an alkaline tolerant photosynthetic bacteria, *R. alkalitolerans*. Since, extremophiles exposed to various environmental factors have the mechanisms to cope with the stresses exposed by the environment, we tried to understand the photoprotective mechanism of this bacteria under various light conditions in relation to its hpH tolerant ability. The absorbance spectroscopy of the isolated intracytoplasmic membranes in alkaline (hpH) condition exhibits a more stable photosystem compared to npH. It’s clearly showing less impact of light stress on the photosystems of the bacteria in hpH than that of the npH (Fig. 2). This result is also complemented by circular dichroism and blue native PAGE of photosynthetic protein supercomplexes. In order to look into the interactions of the pigment-protein, the CD of ICMs was performed at room temperature (RT), wherein the doublet bands at 875 -850 nm show a particular orientational interaction of BChl *a* with that of the scaffold proteins of LH1. Still, the CD pattern in NIR shows almost the same interaction pattern in complexes in three different light conditions (Fig. 3). The CD signal is affected as the culture’s growing in higher light intensities. Still, the degree of reduction was observed from a culture grown in an alkaline medium. It has been shown in the studies that the position of the carotenoid bands as well as the bacteriochlorophyll (BChl) carotenoid ratio affect the NIR- CD spectra (29).

Along with the above results, the content of BChl *a* also has decreased in npH compared to that of the hpH-grown culture (Fig. 1), which indicates that the hpH mitigates the effect of high light. The carotenoid to BChl *a* ratio also has increased in npH compared to hpH (S2). As carotenoid play very important role in photoprotection, by scavenging ROS produced by high light. This signifies that cells grown in hpH are relatively under less photooxidative stress as evident from quantification of level of total ROS and CuZn SOD (Fig. 10). CuZnSOD is also a scavenger of ROS which has been shown to express only in photoheterotrophic growth and detected only when light harvesting complexes are found (20). The growth curve shows the effect of high pH on reaching to stationary phase early compared to npH. This could be because of more utilization of nutrients to produce more energy used in homeostasis and maintaining the intracellular pH. The generation time of bacteria has not shown significant change except in optimum light conditions (Fig. 1, Table.S1). But the hpH-grown cells have entered the stationary phase early compared to that of the npH-grown cells, which indicates that this could be because of the nutrient exhaust where more energy is required for the cell to maintain its cellular homeostasis.

**FIG 10.**
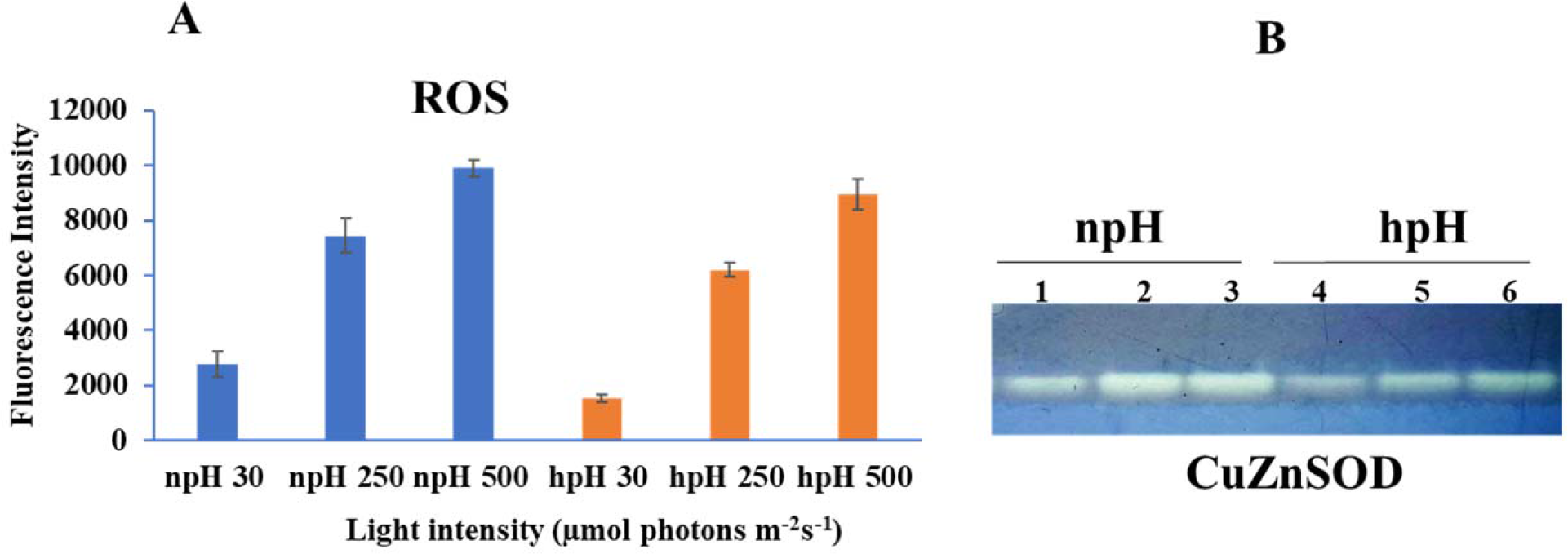
(A) Estimation of total reaction oxygen species (ROS) in cells grown in npH and hpH conditions. (B) Native polyacrylamide gel to detect level of Superoxide dismutase activity in npH and hpH conditions. Lane 1,2,3 represent samples from npH grown at 30,250 and 500 µmol photons m^-2^s^-1^. Likewise lane 4,5,6 sequentially represent sample from hpH grown at 30,250 and 500 µmol photons m^-2^s^-1^.

Measurement of the pH of the culture before harvesting can also be considered as an effect of intracellular metabolism wherein the intracellular pH of the cell in relation to extracellular pH is mainly maintained by antiporters and other metabolites produced by the cell. So, a decrease or increase in the extracellular milieu can result from intracellular machinery. Here the pH of hpH-grown cells has not increased much as the light intensity increase compared to npH-grown cells (Fig. 1). The pH of the culture has increased, indicating the role of antiporters as their expression is obvious in hpH media. As the CD, BN-PAGE, and absorbance spectra show mitigation of light effect, the expression and role of antiporters in light stress mitigation concerning photosynthesis cannot be avoided.

Further, the transmission electron micrograph of cells in each condition strengthens the role of high pH tolerance ability in comparison to normal pH. Fig. 5 shows the effect of light and alkalinity on the morphology of cells and ICMs in different growth conditions wherein the high pH cell ICMs are comparatively not much affected which could be because of the expression of antiporters which get expressed and a similar report has been reported, in other bacteria as well (14). The cell size in the hpH condition is enlarged compared to that of the npH condition, which could be because of delayed assembly of FtsN cell division septum formation as previously identified in *E. coli* when grown in an alkaline condition (21). LP**-**BN- PAGE of photosystem complexes showed five major complexes were identified in each condition when compared with the already published clear native page of *R. sphaeroides* (17). LP-BN shows major changes in RC-LH1 dimeric and monomeric complexes RC-LH1 where the dimeric RC-LH1 complexes have decreased in hpH and monomeric RC-LH1 complexes increased, which could be because of the conversion of dimeric RC-LH1 to monomer, and this pattern is reversed in npH grown cells where both the dimeric and monomeric reaction centers have increased with increase in light intensity. This dimer conversion pattern into monomer is also obvious from the sucrose density gradient, which separates major photosynthetic complexes (Fig. 4,6,7).

Moreover, absorbance spectra of harvested fraction F2 and F3 from sucrose density gradient shows that in hpH the absorption peak at 875nm has increased in F2, which represents the monomeric RC-LH1 with increasing light intensity, but the F3 which represents the dimeric RC-LH1 has decreased. On the contrary to hpH, in npH, the F2 and F3, i.e., the monomeric and dimeric RC-LH1, both have increased in high light. The light-harvesting antenna complex LH2 has decreased with high light in both npH and hpH conditions, although the extent of the light effect is less in hpH (Fig. 4). All these changes were observed because of photoprotection.

Further, more transcript expression levels of subunit-c of ATPase and sodium proton antiporter NhaD in hpH with the increase in light (Fig. 8 b, c). The higher expression would help maintain the homeostasis of the cell by NhaD, and the chromatophore lumen might not be getting more acidified compared to npH- grown cells. Also, the P515 measurement in whole cells in all growth conditions shows that relaxation of the P515 signal is very fast in hpH compared to npH, which signifies the role of ATPase in hpH and its more immediate activity (Fig. 9). As evident from the BN-PAGE and SDG that monomer and dimer both have increased in npH. In contrast, in hpH, the dimer has decreased, and the monomer has increased. This observation also indicates that lumen acidification is essential in forming dimers, and this shows a transition from dimers to monomers is an adaptation strategy. RC-LH1 dimer is also expected to enhance the electron transport rate (22). However, further proton motive force generation would ultimately lead to lumen acidification and become detrimental to photosystem complexes and light-harvesting complexes. Considering the hpH culture condition where the cell must acclimatize to its external milieu of alkaline nature, it requires a lot of protons to maintain its homeostasis, which would be supplied from photosystem machinery, not leading to more acidification of chromatophore lumen and subsequently not much dimerization of RC-LH1 complex. This study also indicates that although pufX is required for the dimerization of the RC-LH1 complex but for, the interaction of two puf X from each monomer of RC- LH1 lumen acidification also would play a role (22).

The role of membrane lipids also could not be avoided in organizing the supercomplexes. The composition of membrane lipids includes Cardiolipin (CL), phosphatidylethanolamine (PE), phosphatidylcholine (PC), phosphatidylglycerol (PG), and sufoquinovocyl diacylglycerol (SQDG) (35). The thin layer chromatographic separation of membrane lipid from the whole cell does not show much difference in PG, PE, CL but SQDG and PC have increased in hpH (Fig. 10 a). PC has been shown to enhance the interaction of pigment-protein complexes and the electron transfer rates between Q_A_ and Q_B_ (18). The role of these membrane lipids in enhancing the supercomplex stability is also evident from CD, blue native PAGE and SDG (Fig. 3 4 5, 6, 7). A case study in *R. sphaeroides* focusing on the role of PC in the stability of the B800-850 complex (23) and chromatophore formation also confirms the importance of this lipid in the current study. At the same time, CL also plays an essential role in photosystem complex stability (35). It might be possible that the conversion of monomer to dimer in the case of hpH- grown cells could be because of a lack of cardiolipin which has slightly decreased in hpH with increased light intensity.

Increase in ROS and SOD levels with increase in light intensity signifies that in photoheterotrophic condition the cells are in more stress as the light intensity increases. As the level of the ROS and SOD that is CuZnSOD are relatively less in hpH condition compared to npH shows that in hpH cells are in relatively less photo-oxidative stress because of an increase in light intensity. CuZnSOD has been shown to express in photo heterotrophic condition confirms that in *Rba. alkalitolerans* there is single type of SOD to express in photoheterotrophic condition (20). Although in related species *R. sphaeroides* FeSOD and CuZnSOD have been reported (20). So comparatively less level of SOD and ROS in hpH might be the reason that the photosystem supercomplexes are relatively stable in hpH. It signifies that in hpH condition *R. alkalitolerans,* till some extent is able to cope up with high light intensity.

Overall, in this study, we tried to investigate the behaviour of photosynthetic apparatus of *R. alkalitolerans* by growing it in alkaline growth condition as it was discovered in an alkaline pond in Gujrat, India. We found that *R. alkalitolerans* cells have comparatively more ICMs and stable photosystem complexes. The level of membrane lipids CL and PC have increased in hpH, which also confers stability to photosystem complexes. Formation of a more dimeric RC-LH1 complex in npH with an increase in light intensity emphasizes importance of its role in acclimatizing as ICM lumen gets more acidified, but ATPase activity is not as efficient as in hpH where dimeric RC-LH1 is converting to monomer along with increased ATPase activity. The reason for dimer to monomer conversion or dimer formation is still elusive, but it can be speculated that lumen acidification might be playing an important role in dimer formation or its interconversion to monomer. Overall, *R. alkalitolerans* cells could adapt to hpH conditions photoheterotrophically with relatively stable photosystem stability and homeostatic balance of the intracellular environment. Relatively less photooxidative stress, higher expression, and activity of ATP synthase from ICMs and Na^+^/H^+^ (NhaD) in hpH in high light indicate the interdependency of homeostasis and photosynthetic machinery functioning in cellular balance inside the cell.

## MATERIALS AND METHODS

### Growth curve and calculation of generation time

For calculating the generation time 8 ml screw-capped tubes were inoculated with 5% of inoculum (v/v) from already grown colony cultures in the log phase. Triplicates of each bacterial culture were taken, and growth in terms of optical density was measured by taking readings at 660 nm (Fig.1, Table 1). Generation time was calculated after plotting the growth curve. Two optical density values were taken from the logarithmic phase of the growth curve, and generation time was calculated as described (24). pH of the culture was measured in all the conditions just before harvesting the cell using Eutech instruments pH tutor.

### Culturing and harvesting of cells

Cultures were grown light/anaerobically as described previously (8) in two pH conditions pH 6.8±.05 and pH 8.6±.05. The cultures were grown in three light intensities of 30 µmol photons m^-2^s^-1^(optimum light), 250-255 µmol photons m^-2^s^-1^, and 500 ± 5 µmol photons m^-2^s^-1^ by inoculating 1.125 ml of inoculum culture in 300 ml of Biebl and Pfenig’s mineral media prepared in 25 mM of Tris buffer media. Cells were harvested in the late log phase by centrifuging the cells at 15,000 x g for 20 min. Washed with 20 mM HEPES pH 7.5 and stored at -80 until required.

### Harvesting of intracytoplasmic membranes

The pellet was washed with 20 mM of HEPES pH 7.5. The washed pellet was again resuspended in 15ml of HEPES buffer and sonicated on ice for 6 min at 30% amplitude by giving 45-second relaxation and 15 min on (25). Before sonication protease inhibitor cocktail (sigma) was also added to avoid protein degradation. Gradients were centrifuged at 50,000×g in a Beckman Type SW32 Ti rotor at 4 °C for 12 h. A pigmented band of ICM formed at the 15/40% interface and was collected using a fixed needle. The membranes were solubilized with 1% n-dodecyl-beta-D-maltoside (β-DM) for 30 min, and non-solubilized material was removed by centrifugation at 10,000x g for 30 min (26,27). The supernatant was loaded in either BN -PAGE or for sucrose density gradient.

### Bacteriochlorophyll *a* (BChl *a*) estimation and carotenoid/BChl *a*

The content of BChl *a* and carotenoid was measured by resuspending the cell pellet in 7 parts of acetone, and 2 parts of methanol (v/v), and absorbance readings were taken at 456 nm and 775 nm, respectively. The content of BChl *a* and carotenoid were calculated as described (28) (Fig. 2). Carotenoid to Bach *a* ratio was also calculated.

### Absorbance Spectroscopy

Absorbance spectra of isolated intracytoplasmic membranes were measured between 400 and 900 nm at room temperature (RT) in a quartz cuvette of 1cm path length at a protein concentration of 50 µg/ml using perkinelmer1500 UV-Visible spectrophotometer (29).

### Sucrose density gradient sedimentation

Sucrose density gradients were formed in transparent ultracentrifuge tubes by carefully layering five steps of 20%, 21.3%, 22.5%, and 23.8%, 25% (w/w) sucrose in 20 mM HEPES (pH 7.5) and 0.03% β-DDM. Solubilized membrane proteins (300 µg) were loaded onto each gradient, and these were centrifuged in a Beckman coulter swing-out bucket rotor SW41Ti at 180,000 x g for 18 h at 4 °C (27). Each strain/growth condition, multiple gradients were run.

### Circular Dichroism spectroscopy

Circular dichroism (CD) spectra were measured at room temperature using a Jasco J-1500 CD spectrometer and a 10 mm path length quartz cuvette with a protein concentration of 50µg/ml. The NIR photomultiplier parameters were set: data pitch: 1 nm, bandwidth: 4 nm, response: 1 s. Two scans for each sample were collected at a speed of 100 nm/min (30), and each sample was in triplicate.

### Large pore Blue Native PAGE (LP-BN)

The first dimension of Blue native gel separation of photosynthetic complexes was carried out in gradient gel of 3.5-12% by solubilization of intracytoplasmic membranes (ICMs) in 1% *n*-dodecyl β-d-maltoside (β-DM) along with the protease inhibitors 1 mM Amino Caproic Acid (ACA), 1 mM benzamidine hydrochloride, the gel with 50 mM ACA is run at 4 °C with a constant current of 4 mA (31,32).

### Identification of subunit protein of super-complexes separated by Blue native -PAGE

Blue- native gel strips were carefully cut and stored at -20 ℃. BN strips were solubilized in lamellae buffer with 6M urea containing β-mercaptoethanol as a reducing agent for 1 h with gentle shaking. Solubilized BN strips were run with 15% SDS-urea gel. Individual protein subunits were spotted, excised, and given for identification of the supercomplexes from BN-PAGE and subsequent quantitative studies in different treatments.

### Transmission electron microscopy (TEM)

To investigate the morphology and number of ICMs in varying light intensities in relation to high pH tolerance, TEM imaging was performed. Harvested cells were washed with potassium phosphate (KPi) buffer, and primary fixation was done with 2% glutaraldehyde (Sigma) for 1 h at 4°C in the dark then cells were washed three times with KPi buffer for 15 min each and post-fixed in 4% KMNO_4_ for 1 h and washed with autoclaved distilled water 5 times each 5 min. After this, cells were fixed with 2% (w/v) uranyl acetate for 1h at room temperature (RT), and after washing with distilled water 5 times each 5 min, cells were subjected to the dehydration step with ethanol (50%,70%,80%,90%, 95% and four washes with 100%. The last two washes were with propylene oxide for 2 times each 1h. These cells were infiltrated with Spurr’s low-viscosity embedding medium, and blocks were prepared. Ultra-thin sections were cut and mounted on copper grids. Sections were stained with an alcoholic solution of 2 % (w/v) uranyl acetate and then with Reynold’s lead citrate stain (33). The thin sections of cells were visualized using a technai instrument (Fig. 5).

### Electrochromic shift measurement

Fast relaxation kinetics measurements were done in Dual PAM-100 (Walz) equipped with P515/P535 emitter- detector module by saturating single turnover flash to intact cells of *R. alkalitolerans.* Cells were incubated in the dark for 1 h before the measurement (34).

### RNA extraction, cDNA synthesis, and quantitative real-time PCR

Total RNA was extracted from STRN50- 1KTspectrum ^TM^ Plant total RNA Kit according to the manufacturer’s protocol. RNA concentration was calculated at 260 nm with a nanodrop1000 spectrophotometer from thermo scientific.

Single-stranded cDNA is synthesized from total RNA by cDNA synthesis kit (TAKARA) by priming at 65 ℃ and reverse transcription for 1 h at 42 ℃ in a 20 µl reaction mixture.

To investigate the expression level of some important genes, primers were designed for ATPase subunit ‘c’ and antiporter NhaD based on the available genome sequence of closely related species *Rhodobacter sphaeroides ATH 2.4.1T (X53853)* (Supplementary Table 1). Using *R. alkalitolerans* cDNA genes were amplified, and 2**^-Δ^ ^ΔCT^** method was used to measure the accurate gene expression of target and housekeeping gene recA. RT-PCR was carried out in Eppendorf Mx3000P multiplex quantitative PCR system with SYBR Green PCR master mix (Kappa).

### Extraction and separation of polar lipids

To extract the polar lipids, 5mg of lyophilized cells were taken. An extraction mixture of methanol-chloroform-water (1:1:0.9, v/v) was added to cells (35) and phase separation was done by centrifuging the mixture at 10,000 rpm for 5 min. The lower layer of chloroform was carefully collected, transferred to another fresh tube, and dried in a speed vacuum. Lipids dried were redissolved in 50 µl of chloroform and kept at -20℃ until use. Total lipid extracts were analysed by TLC on silica gel (20 x 20 cm, layer thickness 0.2 mm). The plate was developed with the solvent chloroform-methanol-acetic acid-water (85: 15:10:3.5, v/v), and detected by iodine vapor.

### Estimation of total ROS and analysis of SOD activity

To estimate the total amount of reactive oxygen species (ROS) cells from all the conditions were collected in logarithmic phase. Fluorescent dye 2′,7′-dichlorofluorescein diacetate (DCFH-DA) (Sigma Aldrich) was used to quantify the total ROS. Cells were washed in culture media and then incubated in the dark by adding 5µM of dye for 1 h at room temperature. After incubation, the cells were washed in culture media to remove excess dye. Fluorescence intensity was measured using a microplate reader (Tecan M250) at excitation wavelength of 485 nm and emission at 530 nm (36).

Also, to analyse the superoxide dismutase activity by native polyacrylamide gel electrophoresis, cells were lysed by sonication, and supernatant was collected. Protein concentration was measured in all the samples and 20 µg of protein was loaded to check the type of superoxide dismutases expressed during the photoheterotrophic growth of bacteria. To visualize the SOD bands, gel was stained in NBT as described previously (37).

## Acknowledgment

RS was supported by the Institute of Eminence (UoH/IoE/RC1/RC1-20-019), DBT-Builder (BT/INF/22/SP41176/2020), DST-FIST and UGC-SAP, Govt. of India, for financial support.

## Author contributions statement

MYZ carried out the experiments, analyzed the data, and drafted the manuscript. RS and CVR designed the experiments, drafted the manuscript, and secured the fund. All authors approved the final version before submission.

## Statement of informed consent, human/animal rights

No conflicts, informed consent, human or animal rights are applicable.

